# Analyzing high-throughput assay data to advance the rapid screening of environmental chemicals for human reproductive toxicity

**DOI:** 10.1101/2024.05.21.595187

**Authors:** Julia R. Varshavsky, Juleen Lam, Courtney Cooper, Patrick Allard, Jennifer Fung, Ashwini Oke, Ravinder Kumar, Joshua F. Robinson, Tracey J. Woodruff

**Affiliations:** Department of Health Sciences and Department of Civil and Environmental Engineering, Northeastern University, Boston, MA; Department of Public Health, California State University, East Bay, Hayward, CA, USA; Institute for Society and Genetics, University of California, Los Angeles (UCLA), Los Angeles, CA, USA; University of California, San Francisco (UCSF), Program on Reproductive Health and the Environment, Department of Obstetrics, Gynecology and Reproductive Sciences, San Francisco, CA, USA; Center for Reproductive Sciences and Department of Obstetrics, Gynecology & Reproductive Sciences, UCSF, San Francisco, CA, USA

**Keywords:** New Approach Methodologies (NAMs), developmental toxicology, chemical assessment, benchmark dose modeling, benchmark response, meiosis, aneuploidy, germ cell development

## Abstract

While high-throughput (HTP) assays have been proposed as platforms to rapidly assess reproductive toxicity, there is currently a lack of established assays that specifically address germline development/function and fertility. We assessed the applicability domains of yeast (*S. cerevisiae)* and nematode *(C. elegans)* HTP assays in toxicity screening of 124 environmental chemicals, determining their agreement in identifying toxicants and their concordance with reproductive toxicity *in vivo*. We integrated data generated in the two models and compared results using a streamlined, semi-automated benchmark dose (BMD) modeling approach. We then extracted and modeled relevant mammalian *in vivo* data available for the matching chemicals included in the Toxicological Reference Database (ToxRefDB). We ranked potencies of common compounds using the BMD and evaluated correlation between the datasets using Pearson and Spearman correlation coefficients. We found moderate to good correlation across the three data sets, with r = 0.48 (95% CI: 0.28–1.00, p<0.001) and r_s_ = 0.40 (p=0.002) for the parametric and rank order correlations between the HTP BMDs; r = 0.95 (95% CI: 0.76–1.00, p=0.0005) and r_s_ = 0.89 (p=0.006) between the yeast assay and ToxRefDB BMDs; and r = 0.81 (95% CI: 0.28–1.00, p=0.014) and r_s_ = 0.75 (p=0.033) between the worm assay and ToxRefDB BMDs. Our findings underscore the potential of these HTP assays to identify environmental chemicals that exhibit reproductive toxicity. Integrating these HTP datasets into mammalian *in vivo* prediction models using machine learning methods could further enhance the predictive value of these assays in future rapid screening efforts.

## 1. Introduction

Difficulty conceiving or carrying a pregnancy to term poses a significant public health burden in the United States, with approximately 10–15% of reproductive-aged couples struggling with fertility problems [1–3]. Multiple adverse reproductive outcomes, including pregnancy loss and birth defects, contribute to this burden of infertility in the U.S. population [4–7] and are associated with impaired germ cell function, including problems with meiosis and early chromosomal aberrations [6,8–11]. Aneuploidy–an abnormal number of chromosomes that occurs from mis-segregation during meiosis–is a leading cause both of miscarriage and major genetic disorders (i.e., Down Syndrome) in humans [12]. Understanding factors that contribute to germ cell disruption could thus alleviate some of the U.S. reproductive health burden.

Although there are more than 80,000 synthetic chemicals in U.S. commerce, the vast majority of have not been adequately tested for potential toxicity to human health [13]. Many of these untested chemicals are similar to those with known toxicities that are suspected to adversely impact fertility and reproductive health. For example, the phenolic compound bisphenol-A (BPA) affects germ cell development, particularly meiosis [14], the key process by which chromosomes segregate during gametogenesis (i.e., when the sperm or oocyte are forming) [8]. Structurally similar compounds, including bisphenol-S (BPS) and certain pesticides, can also have similar effects [15,16]. One critical barrier to understanding the role of environmental chemicals in human reproductive toxicity is studying the influence of chemicals on germ-cell development. Traditional experimental models that test i*n vivo* mammalian toxicity are time consuming and require the use and sacrifice of many animals [17,18]. Subtle effects on germ cells during the earliest stages of fetal development are also difficult to capture through *in vivo* mammalian models [19]. The result is that rodent studies are limited in their ability to rapidly identify germ cell effects, and measured outcomes are sometimes difficult to attribute to direct effects on meiosis or gametogenesis [17,19–21]. This has resulted in scarce information on the reproductive toxicity of environmental contaminants relevant to the human population.

Establishing workflows to rapidly screen environmental chemicals for reproductive toxicity will advance predictive toxicology and chemical assessment. High-throughput (HTP) screening assays can evaluate a broader range of environmental chemicals and incorporate alternative model systems that can test for effects that are typically more difficult to assess or access (i.e., on meiosis and gametogenesis). Updated methods for the rapid screening of environmental chemicals for human reproductive toxicity are required to advance predictive toxicology and risk assessment for the multitude of untested chemicals in commerce. To date, current limitations of HTP assay data have precluded their utility in chemical risk assessment and policy frameworks, with one of the most prominent issues noted being a concern about the limited formal evaluation of existing assays for their specific ability to predict *in vivo* outcomes that are relevant for human health [18]. The inclusion of validated germline assays for reproductive toxicity could thus propel their use in chemical risk assessment frameworks. Moreover, advancing the use of HTP assays would also support broader EPA efforts to integrate new approach methodologies (NAMs) into human health risk assessment and decision-making [22].

To that end, we assessed the performance of two non-mammalian *in vivo* HTP assays that are directly relevant to germline function (i.e., aberrant chromosome segregation or aneuploidy) and fertility: 1) yeast-based (*Saccharomyces cerevisiae) in vivo* assay of meiosis (and mitosis) [23]; and 2) nematode (*Caenorhabditis elegans*) *in vivo* assay which captures chemically-induced meiotic and chromosome errors [24,25]. Previous research has shown that these two assays can predict reproductive toxicity rapidly and cost-effectively. Yeast models are an established platform for screening effects on gametogenesis [26], as demonstrated by research from our collective group, which showed how this model can be used to screen for reproductive toxicants, including BPA and its replacement analogs [23]. Similarly, the *C. elegans* assay has been shown to perform well as a rapid screen for germline toxicants through the induction of aneuploidy [25], a relevant predictor of decreased litter size and ovarian defects in mammalian studies [24].

The objective of this study was to model the dose-response data generated from these two assays for 124 environmental chemicals using a benchmark dose modeling approach and to assess the relationship between germline assays and their ability to predict reproductive toxicity *in vivo*. In this way, we aimed to understand the applicability domain of these models as rapid screens to predict reproductive toxicity.

## 2. Materials and Methods

### 2.1 Project workflow and study design overview

We selected 124 chemicals for rapid screening with our two *in vivo* HTP germline assays based on our previously outlined approach for prioritizing chemicals for toxicity testing [27]. The total list of chemicals, including the CAS numbers and chemical class groupings, can be found in **Supplementary Table S1**. Our project workflow involved evaluating the predictive ability of the two germline/reproductive function assays by first modeling the dose-response data from each HTP assay (yeast and worm) and ranking chemicals from each data set based on potency (**Figure 1**). We further modeled mammalian reproductive dose-response data retrieved from the publicly available Toxicological Reference Database (ToxRefDB) for chemicals that overlapped with our chemical set. ToxRefDB is a comprehensive database of approximately 1,000 chemicals tested in *in vivo* studies [28]. It includes detailed information from published scientific studies, covering aspects such as study design, dosing groups, and observed adverse health endpoints [28]. Notably, it features data on reproductive toxicity endpoints such as ovarian and testicular weight, embryonic and fetal loss, decreased litter size, early postnatal death, and overall fertility decline [28]. Finally, we examined the correlation between all three data sets. Details of each step in the study workflow are described below.

**Figure 1.**
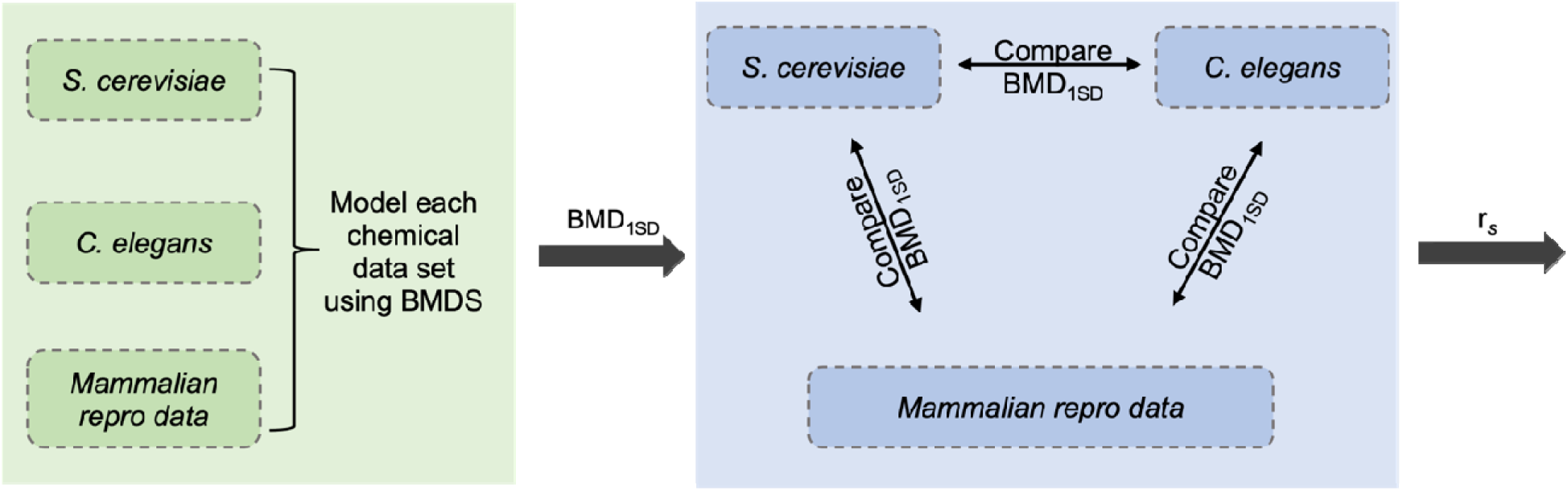
Project workflow for predictive reproductive modeling based on two HTP assays. We modeled dose-response data generated in our two models (*S. cerevisiae and C. elegans)* and available matched *in vivo* datasets from ToxRefDB [28]. Benchmark doses (BMDs) were derived to assess the concordance of toxic potencies between the two HTP assays and their correlation with *in vivo* reproductive toxicity endpoints.

### 2.2 Modeling dose-response data using BMDS software

We used benchmark dose (BMD) modeling to evaluate the dose-response data from the two germ-cell line assays and for the relevant reproductive toxicity tested chemicals in ToxRefDB. BMD modeling is a statistical approach that fits mathematical models to dose-response data to derive an estimate of the dose required to elicit a predefined biological response (for instance, 10% increase in outcome compared to the control group) [29]. BMD modeling incorporates all available data from experimental studies and accounts for the shape of each dataset’s dose-response curve. We used BMDS Wizard (version 1.10) [30] and BMD software (BMDS, version 3.2.1) [31] to automatically calculate BMDs for each chemical following the workflow from a previously published approach [32].

#### 2.2.1 Background on HTP assays (S. cerevisiae and C. elegans)

We used a yeast-based *in vivo* assay to assess multiple doses of each chemical for their potential impacts on meiosis that would lead to aneuploidy (see the companion paper describing the yeast assay by Kumar et al. in this issue). Some of the chemicals were further assessed for the specific stage of meiosis affected and identification of genes and key meiotic processes involved. The yeast assay is based on detecting gamete viability using absorbance measurements to assess the growth of viable gametes through growth curves that shift to the right when the pool of gametes is reduced post exposure [23]. After non-gametes are removed in the assay, the number of viable gametes can be determined by measuring the mitotic growth of living cells that remain in the culture via optical densities at 600 nm (OD_600_) on microplate readers. For this assay, yeast cells were pinned in 96-well plates in three-fold replicates in nitrogen-deprived media to induce meiosis in the absence of chemicals, with meiosis proceeding for 72 hours until full gamete formation. Subsequently, time to half-max – the time to reach halfway point of saturation – is calculated. The shift in time to half-max is used as a proxy for the extent of toxicity for environmental chemicals when comparing to a control such as DMSO (**Supplementary Figure S1A**).

We also used a HTP model system involving an *in vivo* assay for reproductive toxicity and germline impairment in the nematode *C. elegans*, which has been validated with a subset of the EPAs ToxCast phase I library of environmental chemicals and shown to be predictive of mammalian reproductive endpoints [24]. The worm assay can identify chemicals that cause chromosome errors using a fluorescence read-out and can also be paired with other assays to identify which stages of germ cell development are affected. The nematode contains many germ cells, reproduces quickly (with a 4-day life cycle), and is transparent, enabling the practical observation of the reproductive period and allowing for the visualization of germ cell nuclei through all stages of differentiation. The assay involves a fluorescent-based screen called the “Green eggs and HIM” (HIM: High Incidence of Males) of environmental induction of chromosomal errors which can identify errors in germ cell meiotic differentiation that correspond to meiotic segregation errors of the X chromosomes, resulting in the generation of animals carrying only one X chromosome instead of two, and that therefore become male [24,33]. For this assay, 96-well/384-well combination platforms were used, enabling rapid screening of many chemicals in biological quadruplicates. Automated high-content imaging of the plates allowed for high-resolution fluorescence images of the worms in each well. The fluorescent worms were then counted and compared to the total worm count in each well (**Supplementary Figure S1B**).

For both the yeast and the worm assays, DMSO was used as a negative control, while (Latrucilin B) LatB was used as a positive control in the yeast assay and nocodazole was used as a positive control in the worm assay. Each of the HTP assays first tested the response at two dose ranges (30 μM and 100 μM doses) with a subset of 47 chemicals identified as potential “hits” or “positive responses” that were further tested for a wider range of doses (including 10, 50, 125, and 150 μM in the yeast assay and 10, 30, 50, and 100 μM in the worm assay). Other doses were also tested in the subset of chemicals for which we did a more comprehensive dose-response analysis in the *S. cerevisiae* model, including 0.05, 0.10, 0.15, 0.18, 0.30, 0.50, 1.0, 2.0, 5.0, and 60 μM. Laboratory researchers for both assays were blinded to the list of chemicals analyzed, and the chemicals and dosing groups were randomized on well plates to reduce systematic errors. We then modeled the raw yeast assay dose-response data using time to half-max concentration as the response variable on the y-axis and concentration (in μM) as the dose variable on the x-axis. We modeled the raw worm assay data using the positive worm count normalized by total worm count as the continuous response, with DMSO modeled as the control (at 0 μM dose) for both HTP assays. Each of the raw data sets containing dose-response data points (ranging from 0 μM to 150 μM) were exported into a CSV file with individual dose-response data points for use in BMDS modeling.

#### 2.2.2 Background on in vivo mammalian ToxRef datasets

We downloaded publicly available data in June 2018 from EPA’s ToxRef database (ToxRefDB, Version v1.3, released in 2014) [28], selecting studies categorized as the “DevelopmentalReproductive” endpoint category, which included studies with any of the following endpoint targets: organ weight, litter size, viability index, birth index, live birth index, total litter loss, resorptions, live fetuses, reproductive performance, fertility, and dead fetuses. We selected the subset of in vivo studies of reproductive outcomes most relevant to our HTP germline cell assays. We prioritized studies of gonad weight (including ovary and/or testes) as the most sensitive endpoint for our two HTP germline cell assays, based on the decrease of gonad weight observed in many meiotic mutants [17,34–36]. We then identified additional studies with the following relevant reproductive endpoints: viability index, offspring-decrease, litter size – decrease, dead fetuses/live fetuses-decrease, live birth index-decrease, birth index decrease, post implantation loss-increase, aborted pregnancy, NOS-high mortality, decreased number of viable litters, increased pup deaths, postnatal mortality, decrease in 24 hour fetal viability, fertility decrease, implantations-decrease, and resorptions-increase, pregnancy, preimplantation loss, and total litter loss. We did not select studies for HTP germline cell assays including preputial separation, vaginal opening-decrease, delayed descent of testes, and lactation index-decrease, as we deemed these not relevant to our HTP germline cell assays.

The order selection was derived from the functional proximity of these endpoints to the HTP assay, with the top endpoints more likely to reflect issues with germline homeostasis while the bottom endpoints were much more general. We prepared the ToxRef data for BMDS modeling by first extracting the raw data from individual studies, then summarizing the raw data by calculating response means and standard deviations (SDs) for each available dose group.

#### 2.2.3 Benchmark dose modeling parameters and viable model selection

We calculated benchmark doses (BMDs) using BMD Software (BMDS, Version 3.2.1) and a systematic approach to model selection based on previously published decision tree logic [32] consistent with the U.S. EPA’s BMD technical guidance [37]. Briefly, we performed batch calculations of the BMD and its lower limit (BMDL), with a benchmark response (BMR) of 10% extra risk or change in the median equal to 1 SD, for each chemical using the following parameters for each HTP data set: 1) Continuous responses were modeled as either: time to half-max (yeast assay) or the ratio of worms containing at least one positive fluorescent even over the total worm count (*C. elegans* assay); 2) DMSO modeled as the control (0 μM dose) for each HTP data set; 3) BMR type modeled both as a relative difference (BMRF = 0.10) and as a standard deviation (BMRF = 1.0), with the interpretation of the BMD_10_ for a relative difference as follows: the dose (or concentration) at which there is a 10% change in median response relative to the background median (i.e., the median time to half-max or worm count ratio relative to DMSO) and interpretation of the standard deviation BMR type as follows: the dose (or concentration) at which there is a 1 SD change in median relative to the control SD [38]. Models were fit to the dose-response data and assigned to unusable, questionable, and viable categories based on the BMDS Wizard decision logic and prespecified model criteria. Of note, we had to relax the default assumption of constant variance, since the variation in response did not increase or decrease in a monotonic fashion across dose groups in the HTP data sets, which has implications for the interpretation of the BMDL; namely, BMDLs had higher uncertainty and were not stable enough to use reliably as a comparison metric across potency values. Therefore, we used the central estimate (i.e., the BMD) instead as the primary value for potency comparisons and correlation analyses across the three data sets.

Following the Wignall et al. approach [32], we selected viable statistical models from the analyzed dose-response datasets in BMDS based on the following prespecified criteria: if more than one viable model was fit to the data, we selected the model with the lowest Akaike’s information criterion (AIC). If there were no viable models for a particular data set, then we removed the highest dose(s), leaving at least three doses including the control, and remodeled the data in BMDS. We systematically removed the highest dose(s) in each data set if the dose-response curves for a particular data set decreased after first increasing, based on the assumption that a tapering of the dose-response curve would no longer be informative for the point of departure (i.e., BMD) calculation that we were trying to achieve. This was done after we categorized every dose-response curve (with a viable model fit) for each assay based on whether the dose-response curves were increasing, increasing then decreasing, decreasing then increasing, or just decreasing.

### 2.3 Testing correlation between each modeled dose-response dataset

We then evaluated the correlation between derived BMDs from each of the two HTP assays and the correlation between each HTP assay and ToxRef data. We used visual and quantitative tools for the evaluations and comparisons and used parametric statistical tests (i.e., Pearson’s product moment correlation) for log-transformed data (to approximate a normal distribution) and non-parametric statistical tests for distributions that were not normal (i.e., Spearman rank correlation). The tests for significance were calculated under the assumption of a positive association between the data sets for both the Pearson (*R*2) and Spearman (*R*) correlation coefficients (with p-Values < 0.05 considered significant). We removed the *decreasing* and the *decreasing then increasing* dose-response curves prior to the correlation analysis since BMDs from these models would not be informative and would potentially be misleading, for example, by indicating a BMD_1SD_ at which there is a 1 SD change in response even if the change in response is protective (i.e., decreasing). In the decreasing then increasing dose-responses curves, a true decrease in the beginning of the curve was determined based on whether the decrease was statistically significant. All BMD_1SD_ values were log-transformed before the correlation analyses to account for their right-skewed distributions. All correlation analyses were performed in R Studio (Version 2022.07.1).

We selected the most relevant BMD from modeling the ToxRefDB based on our previous prioritization of endpoints (as described in Section 2.2.2): female gonad weight (mg/g), male gonad weight (mg/g), litter size, live fetuses per dam, preimplantation loss, implantations per dam, and resorptions per dam. That is, if a BMD_1SD_ was available from a viable model for female gonad weight, we used that BMD_1SD_ in the correlation analysis. If a viable model for female gonad weight was not available, we used the BMD_1SD_ for male gonad weight. If a viable for male gonad weight was not available, we used the BMD for litter size, and so forth.

### 2.4 Sensitivity analyses

We did a sensitivity analysis for the triclosan results because triclosan had exactly two BMDs to select from our prioritized endpoints. We additionally compared the continuous BMD_1SD_ measurements with categorical determinations of toxicity (i.e., dichotomous “positive response” vs “uncertain” categories) using fold induction (and fold induction / concentration for a “potency” comparison) for the *C. elegans* assay and shift in time to half-max (from DMSO) (and shift in time to half-max / concentration for the potency comparison) for the yeast assay. For the *C. elegans* assay, a positive response was based on whether it was greater than or equal to a pre-defined level of 1.5-fold inductions from DMSO. For the *S. cerevisiae* assay, the pre-defined level was a shift in half-max greater than or equal to 1.5 hours with a p-value cutoff set to 0.05, first at 30 μM then at 100 μM. Chemicals that were considered positive responses at 30 and 100 30 μM were categorized as the most potent, followed by chemicals that were positive responses at 30 μM, with chemicals that were positive responses at 100 μM categorized as the least potent. We further examined positive chemical positive responses in the worm assay as defined by a Z-score value greater than 1.0 at any tested concentration, which included a subset of 13 total chemicals.

Additionally, we compared our BMDS modeling results for the *S. cerevisiae* assay before and after removing dose-response data corresponding to response values that were saturated at half-max of 46 hours, which included 81 data points corresponding to 18 chemicals. Because a half-max response value of 46 represents a manual value assigned when the half-max is beyond the number of hours for which the data is collected (i.e., the maximum possible half-max value above which the assay can no longer accurately determine a response value), the inclusion of saturated response values may introduce bias (because they are right-censored values); therefore, we performed the analysis without these values for comparison.

## 3. Results

### 3.1 Benchmark dose modeling results

Out of a total of N=124 chemicals screened by the HTP assays, we found viable models for n=82 (66%) chemicals with increasing dose-response curves in the yeast assay and n=71 (57%) chemicals in the worm assay, with n=51 (41%) chemicals with viable models overlapping between the yeast and worm assays. Common reasons for questionable or unviable models across chemicals in both assays were: 1) a BMDL 3x lower than the lowest non-zero dose or modeled control response SD > 1.5 of the actual response SD; 2) lack of goodness of fit test calculation or goodness of fit p-value < 0.1; and 3) failure of BMD estimation or computation (due to the lower limit including zero) (**Figure 2**). There were 213 total studies in ToxRef that met our criteria for inclusion, corresponding to 62 chemicals, 51 of which overlapped with our chemical list and 35 of which included gonad weight as an outcome. For this list of 51 chemicals, we were able to locate and download 30 studies, from which we were able to extract raw data on at least one of our outcomes of interest for 24 chemicals. The six studies for which we could not retrieve raw data were missing either the dose, response, and/or standard deviation information needed for BMDS modeling. Of the 24 chemicals from ToxRefDB for which we were able to extract data for BMDS modeling, we found viable models for n=12 chemicals in ToxRefDB; seven ToxRefDB chemicals overlapping with the yeast assay; seven overlapping with the worm assay; and five overlapping across all three data sets (**Figure 2**).

**Figure 2.**
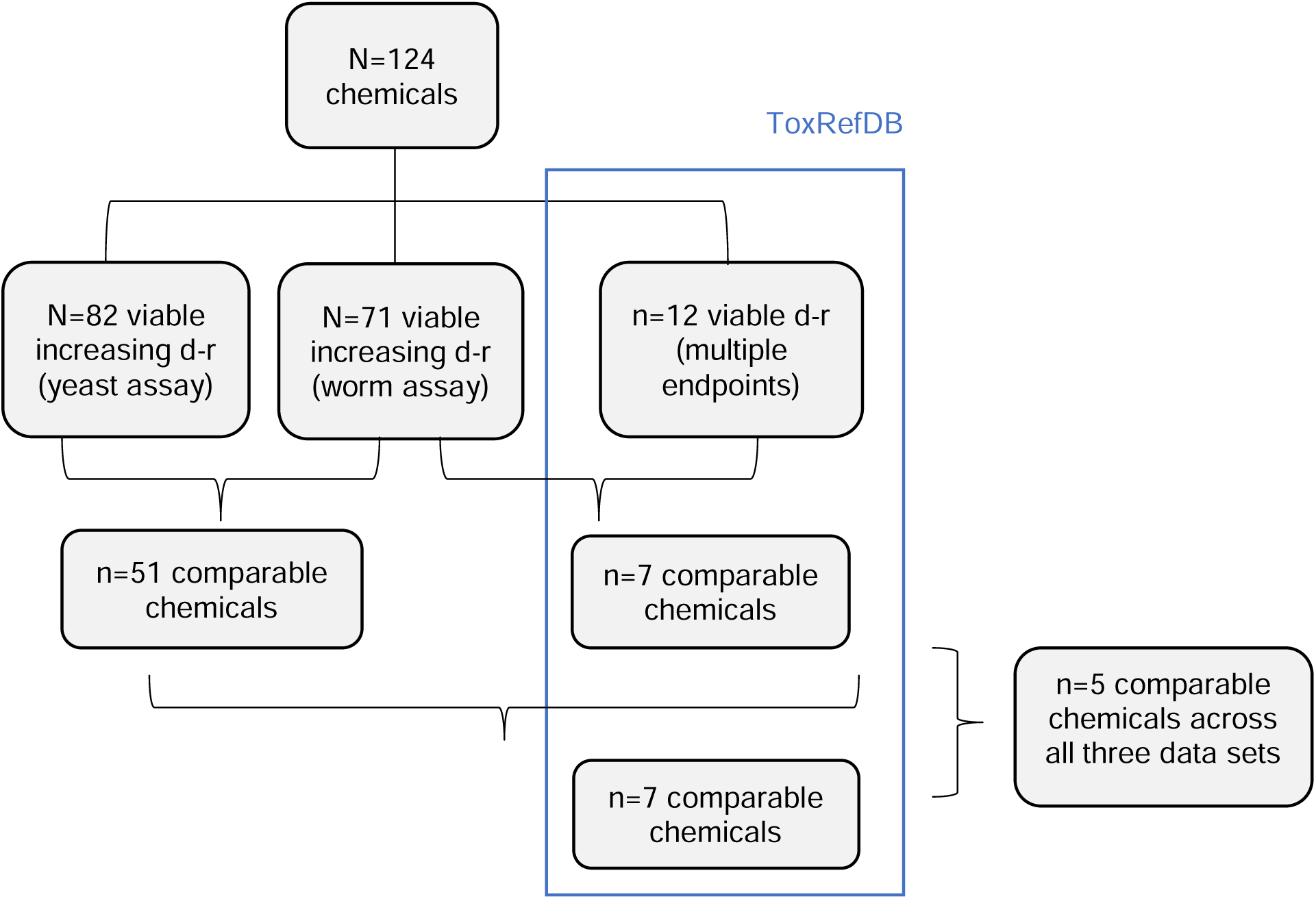
Overview of BMDS modeling results across three data sets (two HTP assays and ToxRefDB).

The BMDs (i.e., BMD_1SD_) for the yeast and worm assays ranged from 0.12–1,353 μM and from 1.82–2,695 μM, respectively (**Table 1 and Supplementary Tables S2 and S3**). The medians were comparable for all the chemicals independent of whether they were the same, with a median of 135 μM in the yeast assay and 145 μM in the worm assay, while the *C. elegans* mean was higher than that of *S. cerevisiae* (with a mean of 234 μM for worm BMD_1SD_ compared to a mean of a mean = 198 μM in the yeast assay). The BMD_1SD_ from the ToxRef database ranged from 32.1–8110, with varying response units based on whether gonad weight (mg/g) rather than a numerical value for litter size or other such endpoint, was modeled in the analysis. Due to the nature of differences in endpoints and units between the HTP assays and ToxRef data, we restricted our BMD models for comparison to the models with benchmark response (BMR) type modeled as the specified change in median relative to the control standard deviation (BMR type set to “Std. Dev.” rather than “Rel. Dev.”) in the BMDS modeling software.

**Table 1.** Variability in BMD_1SD_ between two HTP assay data sets and ToxRefDB.

|  | n | Mean | Median | 25<sup>th</sup> percentile | 75<sup>th</sup> percentile | Range |
| --- | --- | --- | --- | --- | --- | --- |
| *S. cerevisiae* | 82 | 198 | 135 | 30.9 | 253 | 0.12–1353 |
| *C. elegans* | 71 | 234 | 142 | 61.4 | 218 | 1.82–2695 |

Note: dose units were μM for *S. cerevisiae* and *C. elegans* assays and varied in ToxRefDB data from mg/g for gonad weight to number (n) in litter size, live fetuses per dam, preimplantation loss, implantations per dam, and resorptions per dam.

#### 3.1.1 BMD modeling results for yeast assay

Out of the 82 total chemicals with viable increasing dose-response curves from the yeast assay, we observed 13 “more potent” compounds (16% of the total) that were at least one order of magnitude lower than most chemicals, with BMDs that were less than 10 μM (**Figure 3**). There were 17 chemicals (21% of the total) in the middle range (with BMD_1SD_ >10 μM and <100 μM), which was one order of magnitude lower than most chemicals (n=52; 63% of the total) with >100 μM BMD_1SD_ (and two chemicals with BMD_1SD_ above 1000 uM). The most potent 13 chemicals based on BMD_1SD_ modeling (with BMD_1SD_ < 10 μM) included six pesticides (46%), five quaternary ammonium compounds (QACs) (38%), one heavy metal (cadmium chloride) (8%) and one flame retardant (triisopropyl phosphate [TIDPP]) (8%). The pesticides (in order of highest to lowest potency based on lowest to highest BMD_1SD_), were fenbucanazole, triflumizole, fenarimol, spiroxamine, diazinon, and dichlorvos. The QACs (in order of highest to lowest BMD_1SD_ potency) were N, N-Dimethyl-N-benzyl-N-octadecylammonium chloride, Octyl decyl dimethyl ammonium chloride, tetradonium bromide, methylbenzethonium chloride, and benzyldimethyldodecylammonium chloride. The 17 chemicals in the middle range of potency (BMD_1SD_ = 10–100 um) consisted of nine flame retardants (53%), four pesticides (24%), two QACs (12%), and one plastic additive (bisphenol-A [BPA] (6%)). The 52 chemicals in the highest range of potency (BMD_1SD_ > 100 μM) consisted of 16 flame retardants (31%), 15 pesticides (29%), 10 industrial chemicals (19%), six perfluorochemicals (12%), three plasticizers (6%), one pharmaceutical (2%), and one QAC (2%) (**Supplementary Table S2**).

**Figure 3.**
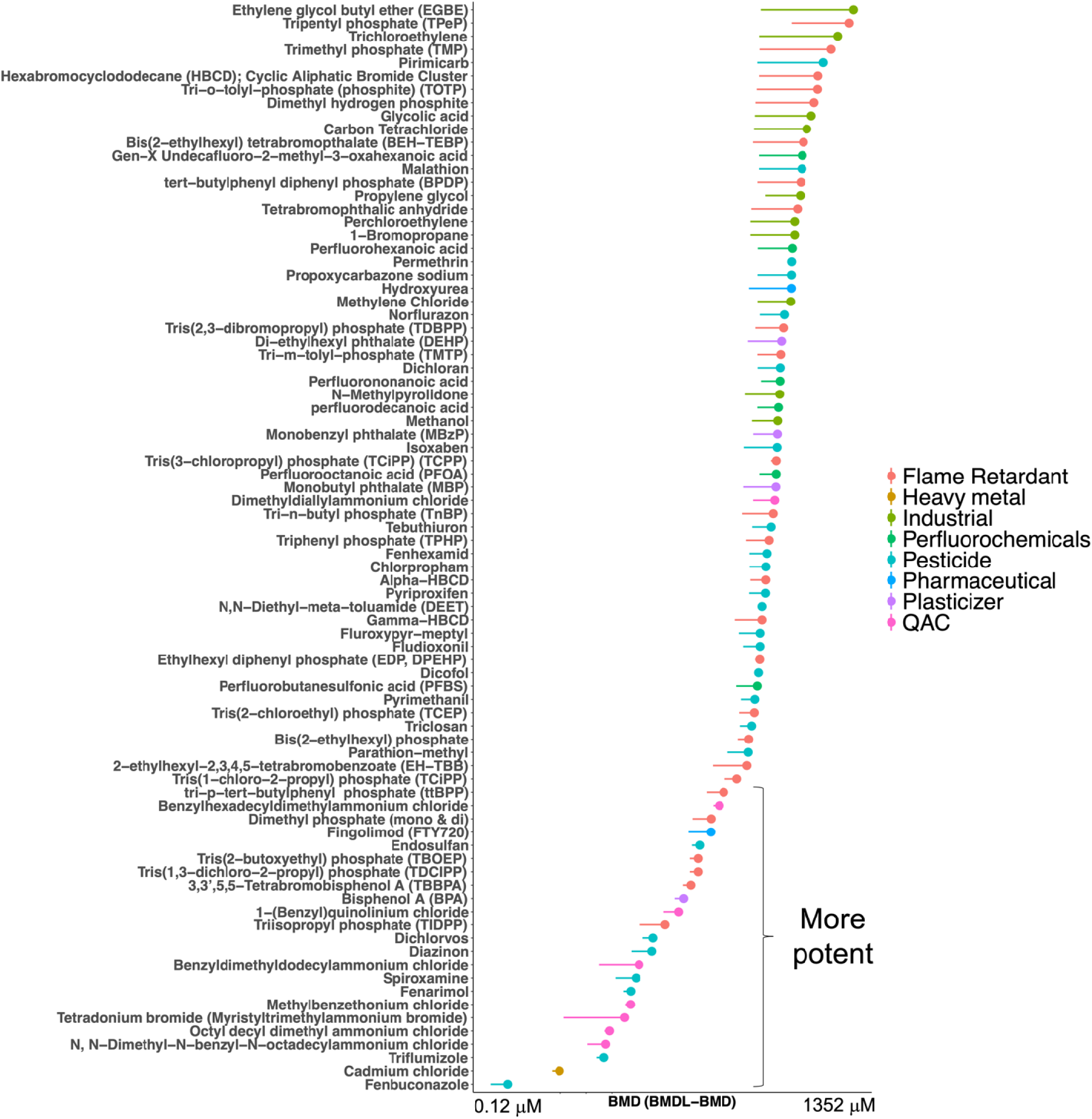
*S. cerevisiae* data BMD_1SD_ modeling results (n=82 chemicals with increasing dose-response curves) color-coded by chemical class.

Eleven out of the 13 chemicals with the lowest BMD (most potent) were also in the highest potency category from the yeast assay (using the >1.5 change in half-max metric). Two chemicals that did not overlap (classified as potent based on BMD_1SD_ models but considered “uncertain” in the yeast assay using the >1.5 change in half-max metric) included the pesticide, diazinon, and the flame retardant, triisopropyl phosphate (TIDPP). Additionally, we did not find a meaningful difference in BMD_1SD_ modeling results from the sensitivity analysis in which we excluded right-censored saturated half-max values. In fact, the exclusion of saturated half-max values in this sensitivity analysis led to less model viability (with viable models achieved for three out of 18 chemicals with saturated half-max response values). For chemicals for which viable models were achieved, the BMD_1SD_ results were not meaningfully different, although one exception was 1-(benzyl)quinolinium chloride), a QAC for which the BMD_1SD_ changed from 12.1 to 8.36, which moved it from the middle potency group (10 uM < BMD_1SD_ < 100 uM) to the higher potency group (BMD_1SD_ < 10 uM).

#### 3.1.2 BMD modeling results for nematode assay

Out of the 71 total chemicals with viable increasing dose-response curves from the worm assay, we observed three “more potent” compounds (4% of the total) that were at least one order of magnitude lower than the majority of chemicals, with BMD_1SD_ that were less than 10 μM (**Figure 4**). There were 24 chemicals (34% of the total) in the middle range (with BMD >10 μM and <100 μM), which was one order of magnitude lower than most chemicals (n=44; 62% of the total) with >100 μM BMD_1SD_ (and three chemicals with BMD_1SD_ above 1000 μM). The most potent three chemicals (with BMDs < 10 μM), in order of lowest to highest BMD_1SD_, included one pesticide (triflumizole) and two QACs (N, N-Dimethyl-N-benzyl-N-octadecylammonium chloride and benzyldimethyldodecylammonium chloride). The 24 chemicals in the middle range of potency (BMD_1SD_ = 10–100 um) consisted of 11 pesticides (17%); three flame retardants (13%); three perfluorochemical (13%); one heavy metal (cadmium chloride) (4%); one pharmaceutical (Acetazolamide) (4%); and one plasticizer (BPA) (4%). The majority of the 44 chemicals in the highest range of potency (BMDs > 100 uM) consisted of 16 flame retardants (36%), 11 pesticides (25%), eight industrial chemicals (18%), three perfluorochemicals (7%), three pharmaceuticals (7%), one plasticizer (2%), one additive (2%), and one QAC (2%) (**Supplementary Table S3**).

**Figure 4.**
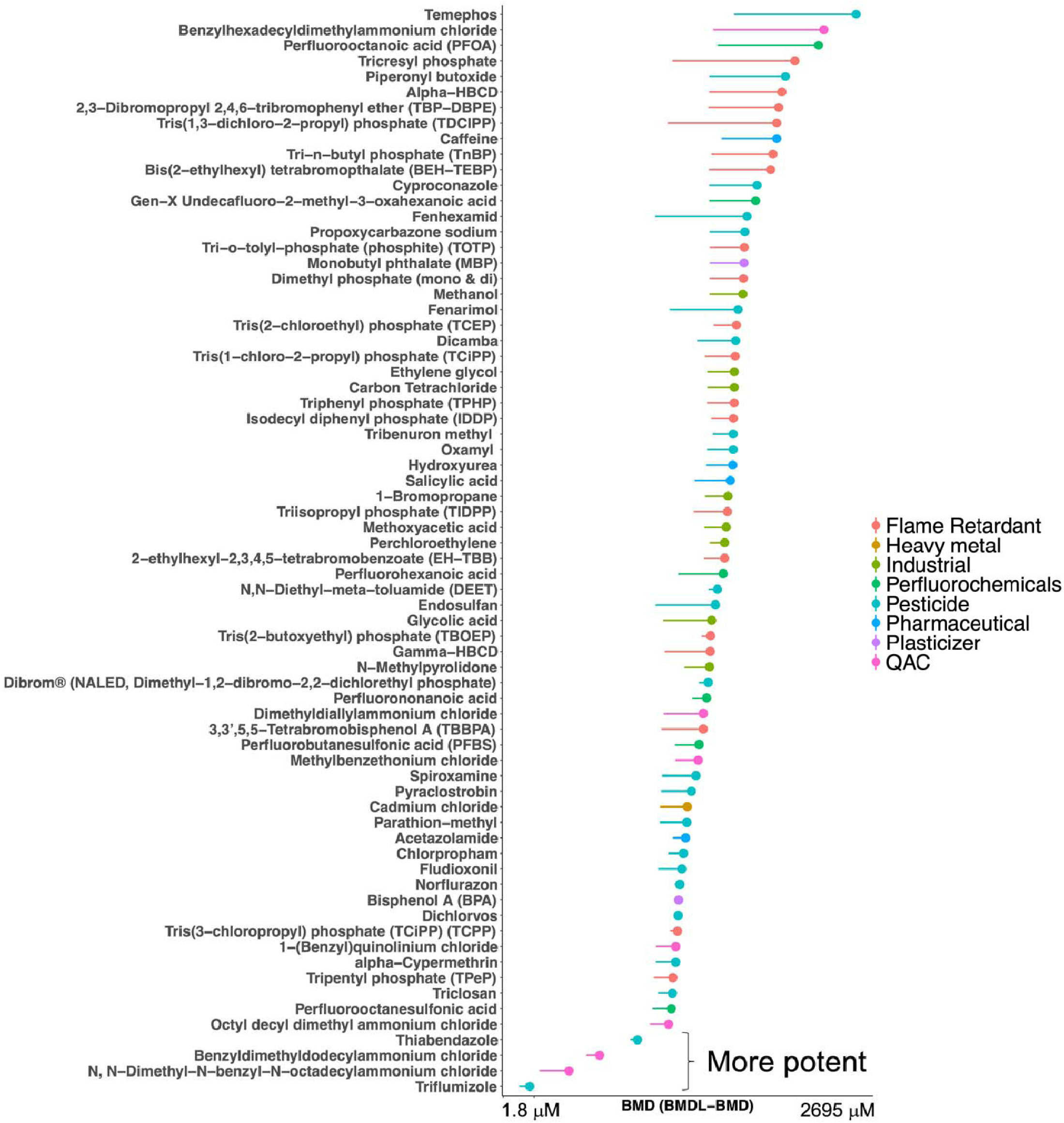
*C. elegans* data BMD_1SD_ modeling results (n=71 chemicals with increasing dose-response curves) color-coded by chemical class.

The three most potent chemicals in the yeast assay (based on BMD-related potency) were also among the five highest potency chemical “positive responses” in the worm assay (which also included methylbenzethonium chloride and fenarimol) based on potency calculations equivalent to the fold induction for each chemical (over DMSO) divided by the dose concentration for each chemical (data not shown). Moreover, those five most potent chemicals were also positive responses in the yeast assay based on a shift in half-max greater than or equal to 1.5 at any concentration. However, DMSO (“DMSO1”) was also included on the worm assay positive response list according to the > 1.5-fold-induction parameter. Therefore, we also compared to the shorter list of worm assay positive responses based on the stricter Z-score > 1 criterion which did not include DMSO1. When incorporating this criterion, fenarimol was not included on the worm assay positive response list, indicating that only four out of five most potent chemicals from BMD_1SD_ modeling were also on the list of positive responses from the worm assay.

#### 3.1.3 BMD modeling results for Toxref data

There were 61 total chemicals with studies from ToxrefDB that met our criteria in evaluating relevant reproductive/developmental endpoint categories (with 27 chemicals for which gonad weight was assessed and 34 for which other reproductive and developmental endpoints were assessed). We found viable models for 12 of the ToxrefDB chemicals (**Figure 2**). Out of those 12 viable models, there were 10 chemicals that overlapped with the list of 124 chemicals that we screened with the HTP assays. For those 10 chemicals, we were able to calculate BMD_1SD_ from viable models corresponding to the endpoints we selected as relevant to our HTP germline cell assays, with viable models for gonad weight achieved for six of the ten chemicals (**Table 2**).

**Table 2.** Benchmark doses (BMD_1SD_)*^a^* for ToxRefDB data based on prioritized endpoint list (from left to right).

| Chemical | Female gonad weight (mg/g) | Male gonad weight (mg/g) | Litter size | Live fetuses per dam | Preimplantation loss | Implantations per dam | Resorptions per dam |
| --- | --- | --- | --- | --- | --- | --- | --- |
| 2,4-Dinitrophenol | **23.1** | Not viable | 70.5 | Not viable | -- | 10.5 | -- |
| BPA | **31.8** | 31.1 | -- | 6.49 | -- | 9.77 | -- |
| DEHP | **785** | 85.7 | -- | 100 | -- | -- | -- |
| TCEP | -- | **280** | -- | -- | -- | -- | -- |
| TCP | -- | **1419** | -- | -- | -- | -- | -- |
| DEET | -- | **605** | -- | -- | -- | -- | -- |
| Malathion | -- | -- | **2079** | 424 | 1477 | 956 | 652 |
| Triclosan | -- | -- | -- | -- | **221** | 496 | -- |
| Pyraclostrobin | -- | -- | -- | **111** | --- | Not viable | 34.9 |
| Hydroxyurea | -- | -- | -- | **462** | -- | -- | -- |

TCEP = Tris(2-chloroethyl) phosphate. Bisphenol-A = BPA. DEET = N,N-Diethyl-meta-toluamide. DEHP = Diethylhexyl phthalate. TCP = Tricresyl phosphate.
<sup>a</sup>Bold color signifies BMD used in correlation analyses. The "--" symbol denotes no data to model whereas "not viable" denotes there was data to model but we were unable to achieve a viable model with the dose-response data. All viable models restricted to F0 generation data only.

### 3.2 Correlation between dose-response data

#### 3.2.1 Concordance of non-mammalian models

We found moderate correlation between the yeast and worm assay BMD data sets (n=51 chemicals available for the comparison), with r = 0.48 (95% CI: 0.28–1.00, p<0.001) for the parametric correlation analysis and r_s_ = 0.40 (p=0.002) for the rank order correlation analysis (**Figure 5**). The QACs, pesticides, industrial chemicals, perfluorochemicals, and pharmaceuticals generally tended to be more highly correlated, and the flame retardants appeared to be less correlated (**Figure 5**). Additionally, using the non-benchmark-based parameters for identifying positive responses in each assay, there were a total of 55 positive responses between the two HTP assays (44 in the worm assay, 31 in the yeast assay, 19 overlapping). When comparing to the narrower list of worm assay positive responses based on Z-score criteria (rather than fold-induction criteria) there were 36 total positive responses between the two assays and eight overlapping positive responses between the two assays.

**Figure 5.**
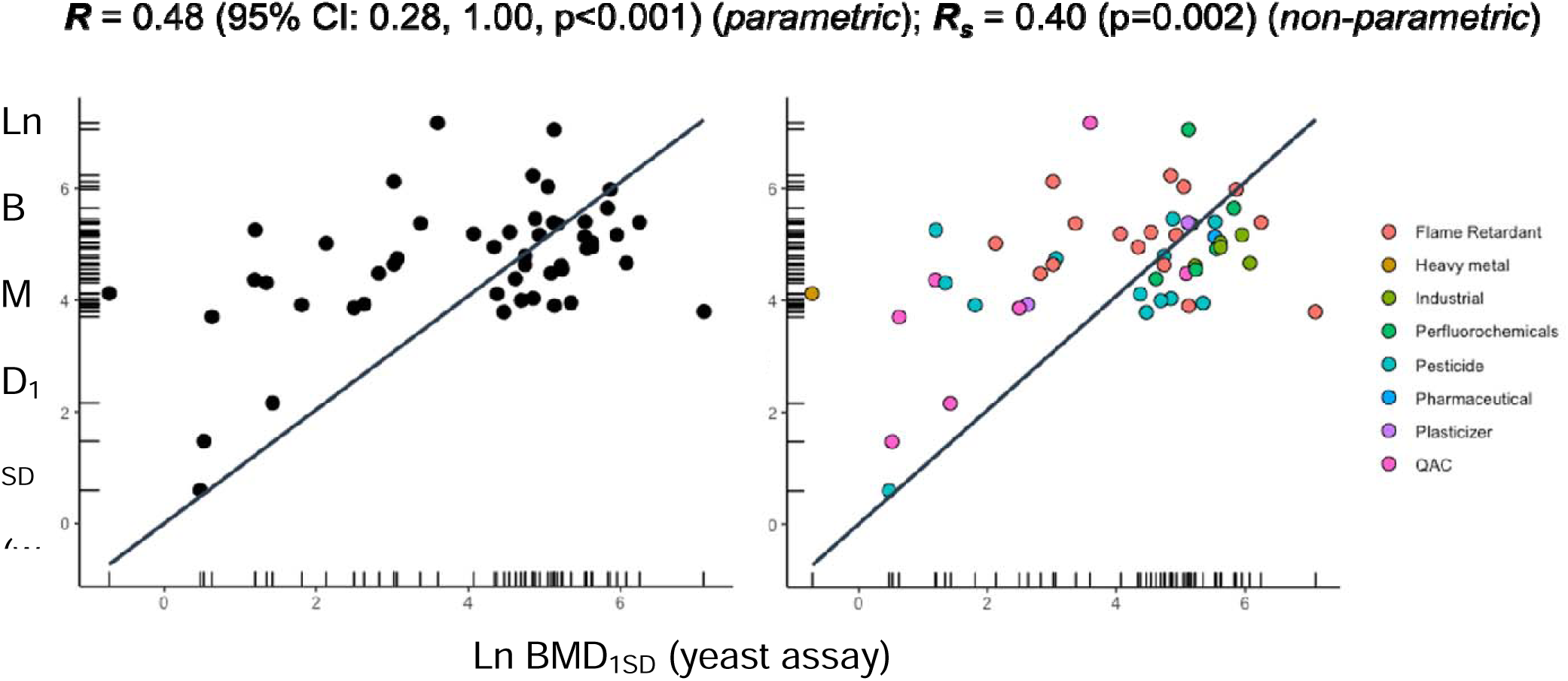
Correlation between *S. cerevisiae* (x-axis) and *C. elegans* (y-axis) BMD_1SD_ data color-coded by chemical class (n=51).

#### 3.2.2 Concordance between yeast or nematode and in vivo data

We found good correlation between BMD_1SD_ from the ToxRefDB and HTP assay data sets, with the Pearson’s correlation coefficients ranging from 0.81 (95% CI: 0.28–1.00, p=0.014) in the worm assay to 0.95 (95% CI: 0.76–1.00, p=0.0005) in the yeast assay, and the spearman’s correlation coefficient ranging from 0.75 (p=0.033) in the worm assay to 0.89 (p=0.006) in the yeast assay (**Figure 6**). When the endpoint prioritization order was changed in our sensitivity analysis from prioritizing preimplantation loss over implantations per dam (to prioritizing implantations per dam over preimplantation loss), the correlation with *S. cerevisiae* data remained relatively constant. However, the correlation to *C. elegans* data was reduced to r = 0.68 (p=0.047) for the Pearson’s product moment correlation and to r*_s_*= 0.43 (p=0.18) for the spearman’s rank correlation (data not shown). The BMD correlation for the five overlapping chemicals between the two HTP assays (BPA, Triclosan, DEET, TCEP, and Hydroxyurea) was less than the correlation between each of the HTP assays and ToxRef data, with the parametric r = 0.65 (95% CI: -0.36–1.00; p=0.113) and the non-parametric r*_s_* = 0.60 (p=0.175).

**Figure 6.**
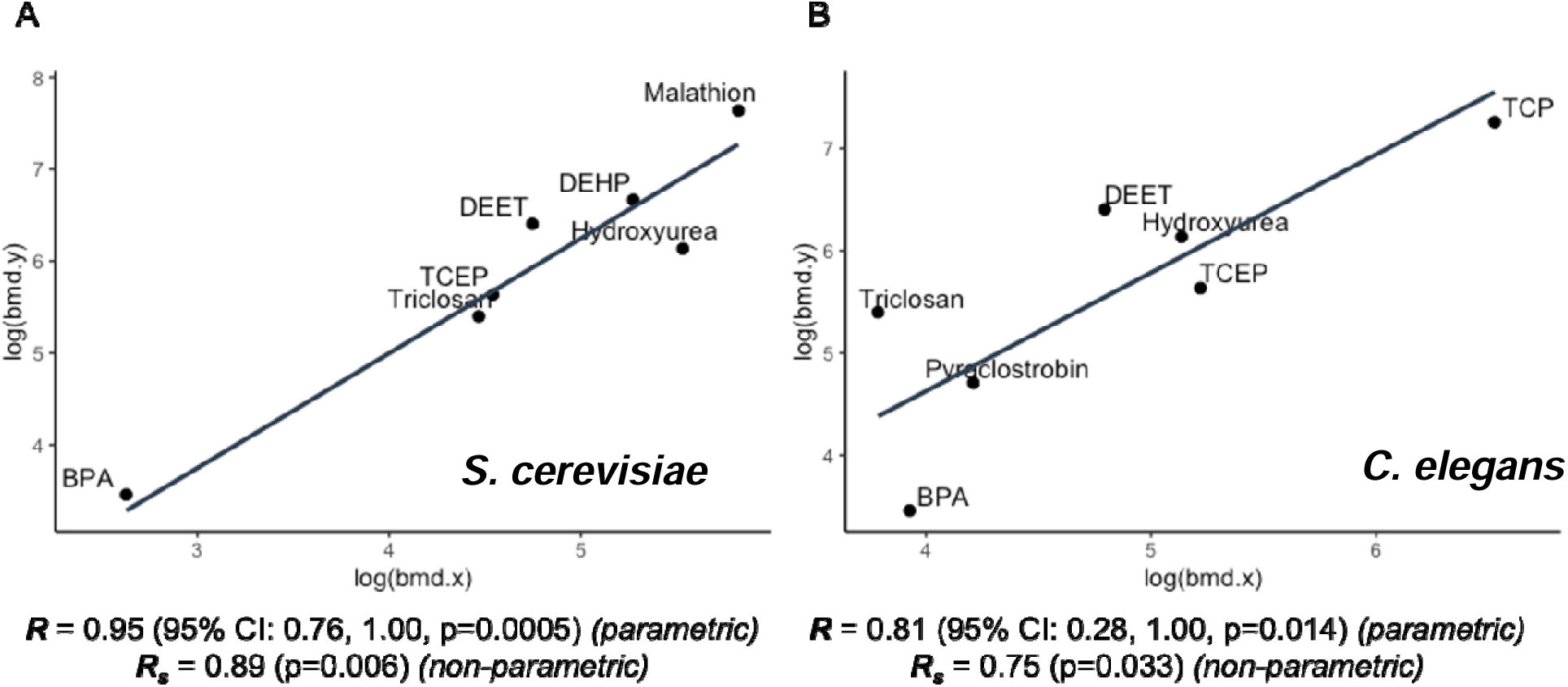
Correlation between the BMD_1SD_ from ToxRef (y-axis) and (A) yeast data (x-axis) or ToxRef (y-axis) and (B) *C. elegans* data (x-axis). There were seven overlapping chemicals in A (BPA, DEET, TCEP, Triclosan, Hydroxyurea, DEHP, and Malathion) and seven overlapping chemicals in B (BPA, DEET, TCEP, Triclosan, Hydroxyurea, Pyraclostrobin, and Tricresyl phosphate [TCP]), with five overlapping chemicals across all three data sets.

## 4. Discussion

This study combined original data generated from two *in vivo* HTP assays with existing publicly available *in vivo* mammalian data to determine the utility of HTP assays for predicting human reproductive toxicity. Specifically, we compared BMDs from the two *in vivo* HTP assays individually and with reproductive findings from mammalian models from EPA’s ToxRefDB database to assess how the assay results compare to each other, how they compare to mammalian *in vivo* data, and to evaluate their predictive ability of the HTP assays for human reproductive toxicity. We found moderate correlation between the HTP data sets and moderate to good correlation between each HTP data set and ToxRefDB. However, there were limited data from which to compare to ToxRefDB, and sensitivity analyses revealed the correlation was dependent on which ToxRefDB endpoints were prioritized for BMD_1SD_ selection. Nevertheless, our results indicate the assays are promising new tools for the rapid screening of environmental chemicals with respect to human reproductive toxicity. Additionally, it’s important to note the limitations with ToxRefDB data in terms of the lack of endpoint specificity regarding germline function, which further highlights the value and importance for using the HTP assays as reproductive screening tools.

### 4.1 Similarities and differences between HTP data sets

We were able to successfully model > 50% of chemicals in both HTP germline cell assays, with several higher potency chemicals identified across the two HTP assays. Pesticides and QACs were the two chemical classes that were most potent in both HTP data sets. Although several of the most potent chemicals in the worm BMD models were also the most potent in the worm assay (i.e., H277; benzyldimethyldodecylammonium chloride), there were other chemicals that ranked high in the worm assay but were not ranked as having higher potency based on our BMD_1SD_ results (i.e., H37; methylbenzethonium chloride). This could in part be due to the shape of the modeling curve, which may or may not reflect the true shape of the dose-response curve for every assayed chemical and modeled dataset. Additionally, we found moderate correlation of the BMD_1SD_ between the HTP assay data sets and between the HTP assay positive responses (as determined without the use of BMD modeling). Some factors that could explain the moderate correlation we observed between HTP data sets include, 1) inherent differences between what the HTP assays are measuring, and 2) the fact that there are many potential sources of variability between HTP data sets (i.e., experimental, endpoint specificity and sensitivity, etc.). For example, norflurazon and diazinon, two chemicals used as positive controls in the worm assay, operate on reproductive toxicity in the worm assay using a different mechanism than in the yeast assay. Moreover, the two assays capture a slightly different set of reproductive endpoints, which also may potentially contribute to divergent results. For example, the worm assay captures reproduction endpoints beyond just meiosis (i.e. chromosomal aberrations in early embryogenesis), while the yeast assay focuses on meiosis (and mitosis) specifically. Other factors that may explain the reason for differing results include experimental differences, such as the differences in dose schemes and use of controls. Further testing with an expanded set of dose groups at the lower range of concentrations (i.e., between 0–30 μM and/or between 30 μM and 100 μM) might be helpful to improve BMD estimation and correlation between the HTP assay data sets going forward. Additional testing within each dose group might also help resolve some of the differences between assay results (e.g., by reducing the variance within each dose group), which could potentially improve the consistency of the BMD_1SD_ values and increase the correlation between assay BMD_1SD_ results.

### 4.2 Correlation with ToxRef data and prediction of mammalian toxicity

In general, we observed good correlation in our comparisons of predicted toxic potencies in yeast or nematode models with *in vivo* data. While these results suggest agreement in conserved response in chemical sensitivity between these non-mammalian vs. rodent models in predicting reproductive toxicity, it is important to exercise caution and consider the limitations in these comparisons which highlight the need for additional data: 1) The results were influenced by the *in vivo* endpoints we considered most sensitive and biologically relevant to the HTP assays; inclusion of different endpoints modified our interpretation in regards to the concordance in HTP vs. *in vivo* results. This has implications for endpoint-specific risk assessment decisions, which may need to rely on a specific assay or endpoints that are more or less sensitive to HTP assay predictions. 2) We were limited in our comparisons due to a lack of available rodent data. While ToxRefDB is a rich resource of publicly available *in vivo* mammalian data, the data which we were able to readily extract and model with BMDS was restricted to seven (yeast) and seven (roundworm) chemicals that overlapped. Extracting additional *in vivo* datasets in the scientific literature maybe an avenue worth exploring to expand this comparison in future studies. Moreover, ToxRefDB studies were conducted more than ten years ago, with the most recent study conducted in 2009 and many studies conducted prior to 2000. Additionally, it is worth noting that potential conflicts of interest within ToxRefDB studies were not evaluated as part of this analysis.

The fact that both HTP assays were individually well correlated with ToxRefDB findings, and that the HTP assay BMDs were also moderately correlated with one another, indicates the assays together may provide a more powerful combination for predicting mammalian reproductive toxicity than they do individually. This makes sense, given the variety of endpoints captured through the combination of assays is more reflective of the diversity of endpoints captured across multiple mammalian *in vivo* studies. A logical next step of this work will be to evaluate the predictive power of both assays with respect to ToxRef data under a machine learning framework. For example, data from both HTP assays in this analysis could be used together, and additionally in combination with EPA’s ToxCast *in vitro* data, to build an unsupervised predictive model of *in vivo* reproductive toxicity. Examining the combined effect of the HTP assays for their collective ability to predict reproductive toxicity will be an important step forward, as the assays are likely to be more informative together than they are alone, given the broader coverage of reproductive endpoints the assays capture together. Future research should also evaluate these assays in combination with other relevant HTP assays, such as assays developed in other species (e.g., zebrafish), which would further advance the use of rapid chemical screens for predicting human reproductive toxicity.

This study integrated dose-response data generated from two *in vivo* rapid HPT germline cell assays combined with publicly available *in vivo* mammalian reproductive toxicity data (on multiple relevant reproductive/developmental endpoints) using computational tools to develop new approaches for assessing potential human health reproductive effects of real-world chemical exposures. More specifically, we compared benchmark dose modeling results across the two model systems (*S. cerevisiae* and *C. elegans*) and in relation to *in vivo* mammalian data from ToxRefDB, which together can be used to better inform potential reproductive and developmental health risks from environmental chemical exposures. We identified pesticides and QACs as potent toxicants in the HTP assays. These findings should be considered in chemical risk assessment and decision-making on these particular compounds. More specifically, our findings should be used to inform efforts to limit the widespread use of QACs, which is a large class of emerging chemicals with antimicrobial and disinfectant properties that has become more common in recent years due to the spread of COVID-19 [39].

Our analysis also sets the stage for future work to integrate and combine HTP datasets in mammalian *in vivo* prediction models using machine learning methods. Further developing rapid screens and integrating new model systems will continue to expand our understanding of reproductive toxicity with respect to human health and should be evaluated together with our HTP assays to further enhance our ability predict human reproductive toxicity using rapid screening assays. Ultimately, our ability to advance rapid screening will provide opportunities to quickly identify which environmental chemical exposures pose the greatest risk for human reproduction and development in sensitive populations, such as pregnant people and their offspring. Thus, our findings will more fully inform evidence-based public health prevention efforts and health-based decision-making to reduce the most harmful chemical exposures and their associated health risks among susceptible populations. This is an important consideration for the EPA as the Agency continues to consider new approach methodologies (NAMs) in their decision-making processes [22].

## Acknowledgements

The authors would like to thank Jeff Gift, US EPA, for technical support and consultation on this project regarding the modeling of dose-response data and use of BMDS software.

## CRediT authorship contribution statement

**Julia Varshavsky:** Data curation, Formal analysis, Investigation, Methodology, Visualization, Writing – original draft. **Juleen Lam:** Conceptualization, Formal analysis, Investigation, Methodology, Supervision, Writing – review & editing. **Courtney Cooper:** Data curation, Project administration, Writing – review & editing. **Patrick Allard:** Data curation, Funding acquisition, Investigation, Writing – review & editing. **Jennifer Fung:** Data curation, Funding acquisition, Investigation, Writing – review & editing. **Ashwini Oke:** Data curation, Writing — review & editing. **Ravinder Kumar**: Investigation. **Joshua Robinson:** Writing – review & editing. **Tracey Woodruff:** Conceptualization, Funding acquisition, Methodology, Supervision, Writing – review & editing.

## Consent for Publication

All the authors gave consent.

---

**Supplementary Figure S1.**
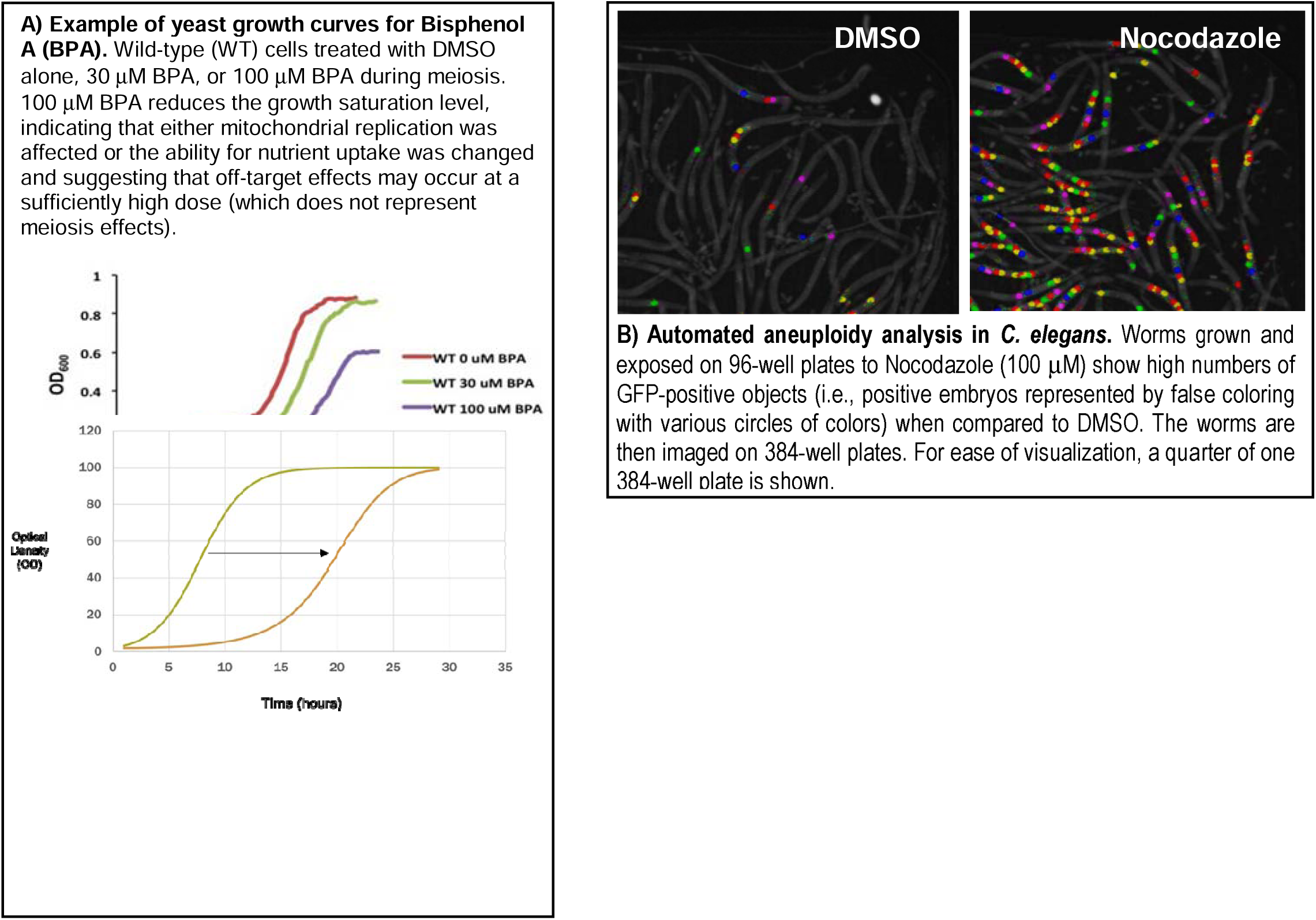
Yeast growth curves (A) and worm counting images (B) from HTP germline assays representing reproductive toxicity.

**Table S1.** Chemical codes, names, and class screened by HTP germline assays (N=124).

| Chemical | CAS # | Name | Class |
| --- | --- | --- | --- |
| 1 | 34014-18-1 | Tebuthiuron | Pesticide |
| 2 | 67-56-1 | Methanol | Industrial |
| 3 | 26444-49-5 | Cresyl diphenyl phosphate (CDP) | Flame Retardant |
| 4 | 26248-87-3 | Tris(3-chloropropyl) phosphate (TCiPP) (TCPP) | Flame Retardant |
| 5 | 183658-27-7 | 2-ethylhexyl-2,3,4,5-tetrabromobenzoate (EH-TBB) | Flame Retardant |
| 6 | 94361-06-5 | Cyproconazole | Pesticide |
| 7 | 13252-13-6 | Gen-X Undecafluoro-2-methyl-3-oxahexanoic acid | Perfluorochemicals |
| 8 | 127-07-1 | Hydroxyurea | Pharmaceutical |
| 9 | 94-26-8 | Butylparaben | Additive |
| 10 | 99-30-9 | Dichloran | Pesticide |
| 11 | 13674-84-5 | Tris(1-chloro-2-propyl) phosphate (TCiPP) | Flame Retardant |
| 12 | 161050-58--4 | Methoxyfenozide | Pesticide |
| 13 | 115-86-6 | Triphenyl phosphate (TPHP) | Flame Retardant |
| 14 | 335-67-1 | Perfluorooctanoic acid (PFOA) | Perfluorochemicals |
| 15 | 115-29-7 | Endosulfan | Pesticide |
| 16 | 335-76-2 | perfluorodecanoic acid | Perfluorochemicals |
| 17 | 868-85-9 | Dimethyl hydrogen phosphite | Flame Retardant |
| 18 | 512-56-1 | Trimethyl phosphate (TMP) | Flame Retardant |
| 19 | 68694-11-1 | Triflumizole | Pesticide |
| 20 | 302-79-4 | Retinoic Acid (all-trans-retinoic acid, RA) | Pharmaceutical |
| 21 | 307-24-4 | Perfluorohexanoic acid | Perfluorochemicals |
| 22 | 112281-77-3 | Tetraconazole | Pesticide |
| 23 | 1119-97-7 | Tetradonium bromide (Myristyltrimethylammonium bromide) | QAC |
| 24 | 95737-68-1 | Pyriproxifen | Pesticide |
| 25 | 298-07-7 | Bis(2-ethylhexyl) phosphate | Flame Retardant |
| 26 | 139-07-1 | Benzyltrimethylammonium chloride | QAC |
| 27 | 3383-96-8 | Temephos | Pesticide |
| 28 | 1241-94-7 | Ethylhexyl diphenyl phosphate (EDP, DPEHP) | Flame Retardant |
| 29 | 78-33-1 | tri-p-tert-butylphenyl phosphate (ttBPP) | Flame Retardant |
| 30 | 333-41-5 | Diazinon | Pesticide |
| 31 | 813-78-5 | Dimethyl phosphate (mono & di) | Flame Retardant |
| 32 | 59-66-5 | Acetazolamide | Pharmaceutical |
| 33 | 122-18-9 | Benzylhexadecyldimethylammonium chloride | QAC |
| 34 | 107-21-1 | Ethylene glycol | Industrial |
| 35 | 52645-53-1 | Permethrin | Pesticide |
| 36 | 298-00-0 | Parathion-methyl | Pesticide |
| 37 | 56803-37-3 | tert-butylphenyl diphenyl phosphate (BPDP) | Flame Retardant |
| 38 | 25155-18-4 | Methylbenzethonium chloride | QAC |
| 39 | 131-70-4 | Monobutyl phthalate (MBP) | Plasticizer |
| 40 | 101-21-3 | Chlorpropham | Pesticide |
| 41 | 175013-18-0 | Pyraclostrobin | Pesticide |
| 42 | 5436-43-1 | 2,2-4,4-Tetrabromodiphenyl ether (BDE-47) | Flame Retardant |
| 43 | 4376-20-9 | Mono-2-Ethylhexyl phthalate (MEHP) | Plasticizer |
| 44 | 872-50-4 | N-Methylpyrrolidone | Industrial |
| 45 | 126833-17-8 | Fenhexamid | Pesticide |
| 46 | 90-43-7 | 2-Phenylphenol | Pesticide |
| 47 | 101200-48-0 | Tribenuron methyl-† | Pesticide |
| 48 | 69-72-7 | Salicylic acid | Pharmaceutical |
| 49 | 148-79-8 | Thiabendazole | Pesticide |
| 50 | 80-05-7 | Bisphenol A (BPA) | Plasticizer |
| 51 | 58-08-2 | Caffeine | Pharmaceutical |
| 52 | 23135-22-0 | Oxamyl-† | Pesticide |
| 53 | 1610-18-0 | Prometon | Pesticide |
| 54 | 106-94-5 | 1-Bromopropane | Industrial |
| 55 | 108-78-1 | Melamine | Flame Retardant |
| 56 | 114369-43-6 | Fenbuconazole | Pesticide |
| 57 | 51-03-6 | Piperonyl butoxide | Pesticide |
| 58 | 131341-86-1 | Fludioxonil | Pesticide |
| 59 | 2528-38-3 | Triphenyl phosphate (TPeP) | Flame Retardant |
| 60 | 121-75-5 | Malathion | Pesticide |
| 61 | 35109-60-5 | 2,3-Dibromopropyl 2,4,6-tribromophenyl ether (TBP-DBPE) | Flame Retardant |
| 62 | 32426-11-2 | Octyl decyl dimethyl ammonium chloride | QAC |
| 63 | 26040-51-7 | Bis(2-ethylhexyl) tetrabromophthalate (BEH-TEBP) | Flame Retardant |
| 64 | 126-72-7 | Tris(2,3-dibromopropyl) phosphate (TDBPP) | Flame Retardant |
| 65 | 79-94-7 | 3,3',5,5'-Tetrabromobisphenol A (TBBPA) | Flame Retardant |
| 66 | 60348-60-9 | 2,2',4,4',5-Penta-bromodiphenyl ether BDE NO. 99 | Flame Retardant |
| 67 | 78-40-0 | Tri-ethyl phosphate (TEP) | Flame Retardant |
| 68 | 23103-98-2 | Pirimicarb | Pesticide |
| 69 | 79-01-6 | Trichloroethylene | Industrial |
| 70 | 53112-28-0 | Pyrimethanil | Pesticide |
| 71 | 134-62-3 | N,N-Diethyl-meta-toluamide (DEET) | Pesticide |
| 72 | 181274-15-7 | Propoxycarbazone sodium | Pesticide |
| 73 | 2528-16-7 | Monobenzyl phthalate (MBzP) | Plasticizer |
| 74 | 81406-37-3 | Fluroxypyr-meptyl | Pesticide |
| 75 | 7173-51-5 | N, N-Dimethyl-N-benzyl-N-octadecylammonium chloride | QAC |
| 76 | 162359-55-9 | Fingolimod (FTY720) | Pharmaceutical |
| 77 | 625-45-6 | Methoxyacetic acid | Industrial |
| 78 | 117-81-7 | Di-ethylhexyl phthalate (DEHP) | Plasticizer |
| 79 | 3380-34-5 | Triclosan | Pesticide |
| 80 | 75-09-2 | Methylene Chloride | Industrial |
| 81 | 1763-23-1 | Perfluorooctanesulfonic acid | Perfluorochemicals |
| 82 | 29761-21-5 | Isodecyl diphenyl phosphate (IDDP) | Flame Retardant |
| 83 | 78-30-8 | Tri-o-tolyl-phosphate (phosphite) (TOTP) | Flame Retardant |
| 84 | 115-96-8 | Tris(2-chloroethyl) phosphate (TCEP) | Flame Retardant |
| 85 | 60168-88-9 | Fenarimol | Pesticide |
| 86 | 563-04-2 | Tri-m-tolyl-phosphate (TMTP) | Flame Retardant |
| 87 | 300-76-5 | Dibrom-Æ (NALED, Dimethyl-1,2-dibromo-2,2-dichlorethyl phosphate) | Pesticide |
| 88 | 57-55-6 | Propylene glycol | Industrial |
| 89 | 15619-48-4 | 1-(Benzyl)quinolinium chloride | QAC |
| 90 | 54-95-5 | D,L-3-hydroxy-3-ethyl-3-phenylpropionamide (HEPP) | Pharmaceutical |
| 91 | 50594-66-6 | Acifluorfen | Pesticide |
| 92 | 128-44-9 | Sodium saccharin | Additive |
| 93 | 375-73-5 | Perfluorobutanesulfonic acid (PFBS) | Perfluorochemicals |
| 94 | 7758-95-4 | Lead Chloride (replaced Lead Acetate) | Heavy metal |
| 95 | 252237-40-4 | Perfluorooctylphosphonic acid | Perfluorochemicals |
| 96 | 27314-13-2 | Norflurazon | Pesticide |
| 97 | 79-14-1 | Glycolic acid | Industrial |
| 98 | 134237-52-8 | Gamma-HBCD | Flame Retardant |
| 99 | 126-73-8 | Tri-n-butyl phosphate (TnBP) | Flame Retardant |
| 100 | 115-32-2 | Dicofol | Pesticide |
| 101 | 111-76-2 | Ethylene glycol butyl ether (EGBE) | Industrial |
| 102 | 632-79-1 | Tetrabromophthalic anhydride | Flame Retardant |
| 103 | 2706-90-3 | Perfluoro-n-pentanoic acid | Perfluorochemicals |
| 104 | 1330-78-5 | Tricresyl phosphate | Flame Retardant |
| 105 | 134237-50-6 | Alpha-HBCD | Flame Retardant |
| 106 | 51-28-5 | 2,4-Dinitrophenol | Phenol |
| 107 | 82558-50-7 | Isoxaben | Pesticide |
| 108 | 3194-55-6 | Hexabromocyclododecane (HBCD); Cyclic Aliphatic Bromide Cluster | Flame Retardant |
| 109 | 1918-00-9 | Dicamba | Pesticide |
| 110 | 134237-51-7 | Beta-HBCD | Flame Retardant |
| 111 | 67375-30-8 | alpha-Cypermethrin | Pesticide |
| 112 | 127-18-4 | Perchloroethylene | Industrial |
| 113 | 7398-69-8 | Dimethyldiallylammonium chloride | QAC |
| 114 | 84087-01-4 | Quinclorac | Pesticide |
| 115 | 513-02-0 | Triisopropyl phosphate (TIDPP) | Flame Retardant |
| 116 | 118134-30-8 | Spiroxamine | Pesticide |
| 117 | 62-73-7 | Dichlorvos | Pesticide |
| 118 | 10108-64-2 | Cadmium chloride | Heavy metal |
| 119 | 22781-23-3 | Bendiocarb | Pesticide |
| 120 | 78-51-3 | Tris(2-butoxyethyl) phosphate (TBOEP) | Flame Retardant |
| 121 | 375-95-1 | Perfluorononanoic acid | Perfluorochemicals |
| 122 | 56-23-5 | Carbon Tetrachloride | Industrial |
| 123 | 13674-87-8 | Tris(1,3-dichloro-2-propyl) phosphate (TDCIPP) | Flame Retardant |
| 124 | 375-85-9 | Perfluoroheptanoic acid | Perfluorochemicals |

**Table S2.** Ordered list of BMD1SD and BMDL for chemicals with viable models in the *S. cerevisiae* germline assay (N=82).

| CAS # | Chemical | BMD | BMDL | Class |
| --- | --- | --- | --- | --- |
| 111-76-2 | Ethylene glycol butyl ether (EGBE) | 1353.37 | 111.57 | Industrial |
| 2528-38-3 | Triphenyl phosphate (TPeP) | 1203.83 | 254.59 | Flame Retardant |
| 79-01-6 | Trichloroethylene | 888.64 | 106.93 | Industrial |
| 512-56-1 | Trimethyl phosphate (TMP) | 741.86 | 108.00 | Flame Retardant |
| 23103-98-2 | Pirimicarb | 601.74 | 101.43 | Pesticide |
| 3194-55-6 | Hexabromocyclododecane (HBCD); Cyclic Aliphatic Bromide Cluster | 521.16 | 106.45 | Flame Retardant |
| 78-30-8 | Tri-o-tolyl-phosphate (phosphite) (TOTP) | 517.35 | 99.85 | Flame Retardant |
| 868-85-9 | Dimethyl hydrogen phosphite | 469.15 | 96.47 | Flame Retardant |
| 79-14-1 | Glycolic acid | 434.69 | 94.93 | Industrial |
| 56-23-5 | Carbon Tetrachloride | 384.53 | 92.71 | Industrial |
| 26040-51-7 | Bis(2-ethylhexyl) tetrabromophthalate (BEH-TEBP) | 351.93 | 90.19 | Flame Retardant |
| 13252-13-6 | Gen-X Undecafluoro-2-methyl-3-oxahexanoic acid | 340.33 | 106.43 | Perfluorochemicals |
| 121-75-5 | Malathion | 337.08 | 106.33 | Pesticide |
| 56803-37-3 | tert-butylphenyl diphenyl phosphate (BPDP) | 328.30 | 101.77 | Flame Retardant |
| 57-55-6 | Propylene glycol | 323.96 | 125.67 | Industrial |
| 632-79-1 | Tetrabromophthalic anhydride | 299.69 | 86.21 | Flame Retardant |
| 127-18-4 | Perchloroethylene | 276.68 | 84.16 | Industrial |
| 106-94-5 | 1-Bromopropane | 276.23 | 84.11 | Industrial |
| 307-24-4 | Perfluorohexanoic acid | 259.19 | 102.82 | Perfluorochemicals |
| 52645-53-1 | Permethrin | 254.91 | 233.60 | Pesticide |
| 181274-15-7 | Propoxycarbazon sodium | 253.64 | 102.07 | Pesticide |
| 127-07-1 | Hydroxyurea | 252.40 | 80.96 | Pharmaceutical |
| 75-09-2 | Methylene Chloride | 248.33 | 102.13 | Industrial |
| 27314-13-2 | Norflurazon | 210.13 | 108.70 | Pesticide |
| 126-72-7 | Tris(2,3-dibromopropyl) phosphate (TDBPP) | 205.00 | 96.20 | Flame Retardant |
| 117-81-7 | Di-ethylhexyl phthalate (DEHP) | 194.91 | 78.44 | Plasticizer |
| 563-04-2 | Tri-m-tolyl-phosphate (TMTP) | 190.45 | 101.62 | Flame Retardant |
| 99-30-9 | Dichloran | 187.85 | 102.40 | Pesticide |
| 375-95-1 | Perfluorononanoic acid | 187.02 | 112.28 | Perfluorochemicals |
| 872-50-4 | N-Methylpyrrolidone | 185.16 | 72.89 | Industrial |
| 335-76-2 | perfluorodecanoic acid | 178.74 | 101.52 | Perfluorochemicals |
| 67-56-1 | Methanol | 175.38 | 87.82 | Industrial |
| 2528-16-7 | Monobenzyl phthalate (MBzP) | 174.12 | 90.84 | Plasticizer |
| 82558-50-7 | Isoxaben | 172.58 | 70.72 | Pesticide |
| 26248-87-3 | Tris(3-chloropropyl) phosphate (TCiPP) (TCPP) | 168.79 | 146.01 | Flame Retardant |
| 335-67-1 | Perfluorooctanoic acid (PFOA) | 168.13 | 107.58 | Perfluorochemicals |
| 131-70-4 | Monobutyl phthalate (MBP) | 166.93 | 69.77 | Plasticizer |
| 7398-69-8 | Dimethyldiallylammonium chloride | 161.83 | 90.54 | QAC |
| 126-73-8 | Tri-n-butyl phosphate (TnBP) | 155.24 | 67.55 | Flame Retardant |
| 34014-18-1 | Tebuthiuron | 146.70 | 88.40 | Pesticide |
| 115-86-6 | Triphenyl phosphate (TPHP) | 138.89 | 74.88 | Flame Retardant |
| 126833-17-8 | Fenhexamid | 131.19 | 81.94 | Pesticide |
| 101-21-3 | Chlorpropham | 127.66 | 82.18 | Pesticide |
| 134237-50-6 | Alpha-HBCD | 127.14 | 84.00 | Flame Retardant |
| 95737-68-1 | Pyriproxifen | 126.27 | 81.35 | Pesticide |
| 134-62-3 | N,N-Diethyl-meta-toluamide (DEET) | 115.18 | 104.02 | Pesticide |
| 134237-52-8 | Gamma-HBCD | 115.12 | 55.51 | Flame Retardant |
| 81406-37-3 | Fluroxypyr-meptyl | 109.44 | 61.97 | Pesticide |
| 131341-86-1 | Fludioxonil | 109.27 | 69.53 | Pesticide |
| 1241-94-7 | Ethylhexyl diphenyl phosphate (EDP, DPEHP) | 108.46 | 95.62 | Flame Retardant |
| 115-32-2 | Dicofol | 104.86 | 97.67 | Pesticide |
| 375-73-5 | Perfluorobutanesulfonic acid (PFBS) | 101.41 | 57.67 | Perfluorochemicals |
| 53112-28-0 | Pyrimethanil | 94.86 | 65.61 | Pesticide |
| 115-96-8 | Tris(2-chloroethyl) phosphate (TCEP) | 93.74 | 62.47 | Flame Retardant |
| 3380-34-5 | Triclosan | 87.14 | 63.69 | Pesticide |
| 298-07-7 | Bis(2-ethylhexyl) phosphate | 80.47 | 60.25 | Flame Retardant |
| 298-00-0 | Parathion-methyl | 79.38 | 45.43 | Pesticide |
| 183658-27-7 | 2-ethylhexyl-2,3,4,5-tetrabromobenzoate (EH-TBB) | 76.69 | 30.38 | Flame Retardant |
| 13674-84-5 | Tris(1-chloro-2-propyl) phosphate (TCiPP) | 58.43 | 42.04 | Flame Retardant |
| 78-33-1 | tri-p-tert-butylphenyl phosphate (ttBPP) | 40.89 | 25.79 | Flame Retardant |
| 122-18-9 | Benzylhexadecyldimethylammonium chloride | 36.43 | 30.77 | QAC |
| 813-78-5 | Dimethyl phosphate (mono & di) | 29.03 | 17.60 | Flame Retardant |
| 162359-55-9 | Fingolimod (FTY720) | 28.80 | 15.78 | Pharmaceutical |
| 115-29-7 | Endosulfan | 21.38 | 17.32 | Pesticide |
| 78-51-3 | Tris(2-butoxyethyl) phosphate (TBOEP) | 20.49 | 16.33 | Flame Retardant |
| 13674-87-8 | Tris(1,3-dichloro-2-propyl) phosphate (TDCIPP) | 20.49 | 16.36 | Flame Retardant |
| 79-94-7 | 3,3',5,5-Tetrabromobisphenol A (TBBPA) | 16.85 | 13.59 | Flame Retardant |
| 80-05-7 | Bisphenol A (BPA) | 13.87 | 10.91 | Plasticizer |
| 15619-48-4 | 1-(Benzyl)quinolinium chloride | 12.14 | 8.07 | QAC |
| 513-02-0 | Triisopropyl phosphate (TIDPP) | 8.41 | 4.18 | Flame Retardant |
| 62-73-7 | Dichlorvos | 6.10 | 4.61 | Pesticide |
| 333-41-5 | Diazinon | 5.91 | 3.38 | Pesticide |
| 139-07-1 | Benzyltrimethylammonium chloride | 4.16 | 1.41 | QAC |
| 118134-30-8 | Spiroamine | 3.84 | 2.20 | Pesticide |
| 60168-88-9 | Fenarimol | 3.32 | 2.72 | Pesticide |
| 25155-18-4 | Methylbenzethonium chloride | 3.29 | 2.85 | QAC |
| 1119-97-7 | Tetradonium bromide (Myristyltrimethylammonium bromide) | 2.80 | 0.55 | QAC |
| 32426-11-2 | Octyl decyl dimethyl ammonium chloride | 1.87 | 1.64 | QAC |
| 7173-51-5 | N, N-Dimethyl-N-benzyl-N-octadecylammonium chloride | 1.69 | 1.03 | QAC |
| 68694-11-1 | Triflumizole | 1.61 | 1.33 | Pesticide |
| 10108-64-2 | Cadmium chloride | 0.49 | 0.40 | Heavy metal |
| 114369-43-6 | Fenbuconazole | 0.12 | 0.08 | Pesticide |

**Table S3.** Ordered list of BMD1SD and BMDL for chemicals with viable models in the *C. elegans* germline assay (N=71).

| CAS # | Chemical | BMD | BMDL | Class |
| --- | --- | --- | --- | --- |
| 3383-96-8 | Temephos | 2695.36 | 174.94 | Pesticide |
| 122-18-9 | Benzylhexadecyldimethylammonium chloride | 1306.92 | 109.81 | QAC |
| 335-67-1 | Perfluorooctanoic acid (PFOA) | 1160.78 | 121.46 | Perfluorochemicals |
| 1330-78-5 | Tricresyl phosphate | 683.12 | 44.05 | Flame Retardant |
| 51-03-6 | Piperonyl butoxide | 553.96 | 100.40 | Pesticide |
| 134237-50-6 | Alpha-HBCD | 510.01 | 100.40 | Flame Retardant |
| 35109-60-5 | 2,3-Dibromopropyl 2,4,6-tribromophenyl ether (TBP-DBPE) | 476.79 | 98.90 | Flame Retardant |
| 128-44-9 | Sodium saccharin | 460.68 | 55.48 | Additive |
| 13674-87-8 | Tris(1,3-dichloro-2-propyl) phosphate (TDCIPP) | 456.42 | 39.53 | Flame Retardant |
| 58-08-2 | Caffeine | 455.94 | 132.58 | Pharmaceutical |
| 126-73-8 | Tri-n-butyl phosphate (TnBP) | 418.07 | 105.46 | Flame Retardant |
| 26040-51-7 | Bis(2-ethylhexyl) tetrabromophthalate (BEH-TEBP) | 397.77 | 99.61 | Flame Retardant |
| 94361-06-5 | Cyproconazole | 293.33 | 100.25 | Pesticide |
| 13252-13-6 | Gen-X Undecafluoro-2-methyl-3-oxahexanoic acid | 284.18 | 100.54 | Perfluorochemicals |
| 126833-17-8 | Fenhexamid | 234.75 | 30.01 | Pesticide |
| 181274-15-7 | Propoxycarbazone sodium | 222.77 | 101.48 | Pesticide |
| 78-30-8 | Tri-o-tolyl-phosphate (phosphite) (TOTP) | 220.51 | 101.85 | Flame Retardant |
| 131-70-4 | Monobutyl phthalate (MBP) | 219.25 | 101.73 | Plasticizer |
| 813-78-5 | Dimethyl phosphate (mono & di) | 216.35 | 101.43 | Flame Retardant |
| 67-56-1 | Methanol | 213.31 | 101.11 | Industrial |
| 60168-88-9 | Fenarimol | 192.03 | 41.66 | Pesticide |
| 115-96-8 | Tris(2-chloroethyl) phosphate (TCEP) | 184.89 | 111.31 | Flame Retardant |
| 1918-00-9 | Dicamba | 181.82 | 76.58 | Pesticide |
| 13674-84-5 | Tris(1-chloro-2-propyl) phosphate (TCIPP) | 179.53 | 91.51 | Flame Retardant |
| 107-21-1 | Ethylene glycol | 176.35 | 96.20 | Industrial |
| 56-23-5 | Carbon Tetrachloride | 176.00 | 96.15 | Industrial |
| 115-86-6 | Triphenyl phosphate (TPHP) | 175.62 | 95.54 | Flame Retardant |
| 29761-21-5 | Isodecyl diphenyl phosphate (IDDP) | 172.67 | 105.39 | Flame Retardant |
| 101200-48-0 | Tribenuron methyl-† | 171.98 | 108.70 | Pesticide |
| 23135-22-0 | Oxamyl-† | 171.83 | 95.44 | Pesticide |
| 127-07-1 | Hydroxyurea | 170.04 | 94.29 | Pharmaceutical |
| 69-72-7 | Salicylic acid | 161.48 | 72.66 | Pharmaceutical |
| 106-94-5 | 1-Bromopropane | 153.14 | 91.75 | Industrial |
| 513-02-0 | Triisopropyl phosphate (TIDPP) | 151.50 | 71.15 | Flame Retardant |
| 625-45-6 | Methoxyacetic acid | 147.23 | 90.41 | Industrial |
| 127-18-4 | Perchloroethylene | 141.82 | 101.68 | Industrial |
| 183658-27-7 | 2-ethylhexyl-2,3,4,5-tetrabromobenzoate (EH-TBB) | 141.69 | 89.14 | Flame Retardant |
| 307-24-4 | Perfluorohexanoic acid | 137.35 | 50.11 | Perfluorochemicals |
| 134-62-3 | N,N-Diethyl-meta-toluamide (DEET) | 120.88 | 98.99 | Pesticide |
| 115-29-7 | Endosulfan | 115.45 | 30.23 | Pesticide |
| 79-14-1 | Glycolic acid | 106.29 | 35.87 | Industrial |
| 78-51-3 | Tris(2-butoxyethyl) phosphate (TBOEP) | 103.72 | 84.71 | Flame Retardant |
| 134237-52-8 | Gamma-HBCD | 102.87 | 37.10 | Flame Retardant |
| 872-50-4 | N-Methylpyrrolidone | 101.41 | 57.70 | Industrial |
| 300-76-5 | Dibrom-Æ (NALED, Dimethyl-1,2-dibromo-2,2-dichlorethyl phosphate) | 98.44 | 79.72 | Pesticide |
| 375-95-1 | Perfluorononanoic acid | 94.81 | 68.93 | Perfluorochemicals |
| 7398-69-8 | Dimethyldiallylammonium chloride | 88.13 | 36.36 | QAC |
| 79-94-7 | 3,3',5,5'-Tetrabromobisphenol A (TBBPA) | 88.07 | 34.49 | Flame Retardant |
| 375-73-5 | Perfluorobutanesulfonic acid (PFBS) | 80.20 | 46.87 | Perfluorochemicals |
| 25155-18-4 | Methylbenzethonium chloride | 78.69 | 47.41 | QAC |
| 118134-30-8 | Spiroxamine | 74.58 | 35.02 | Pesticide |
| 175013-18-0 | Pyraclostrobin | 67.40 | 34.29 | Pesticide |
| 10108-64-2 | Cadmium chloride | 61.69 | 33.50 | Heavy metal |
| 298-00-0 | Parathion-methyl | 61.01 | 33.31 | Pesticide |
| 59-66-5 | Acetazolamide | 59.25 | 44.58 | Pharmaceutical |
| 101-21-3 | Chlorpropham | 56.45 | 40.18 | Pesticide |
| 131341-86-1 | Fludioxonil | 54.44 | 32.11 | Pesticide |
| 27314-13-2 | Norflurazon | 52.11 | 46.42 | Pesticide |
| 80-05-7 | Bisphenol A (BPA) | 50.77 | 49.24 | Plasticizer |
| 62-73-7 | Dichlorvos | 50.18 | 48.93 | Pesticide |
| 26248-87-3 | Tris(3-chloropropyl) phosphate (TCiPP) (TCPP) | 49.42 | 41.89 | Flame Retardant |
| 15619-48-4 | 1-(Benzyl)quinolinium chloride | 47.51 | 30.51 | QAC |
| 67375-30-8 | alpha-Cypermethrin | 47.48 | 30.47 | Pesticide |
| 2528-38-3 | Triphenyl phosphate (TPeP) | 44.56 | 29.18 | Flame Retardant |
| 3380-34-5 | Triclosan | 44.13 | 32.02 | Pesticide |
| 1763-23-1 | Perfluorooctanesulfonic acid | 43.04 | 28.20 | Perfluorochemicals |
| 32426-11-2 | Octyl decyl dimethyl ammonium chloride | 40.57 | 26.70 | QAC |
| 148-79-8 | Thiabendazole | 20.24 | 17.23 | Pesticide |
| 139-07-1 | Benzyltrimethylammonium chloride | 8.66 | 6.42 | QAC |
| 7173-51-5 | N, N-Dimethyl-N-benzyl-N-octadecylammonium chloride | 4.37 | 2.26 | QAC |
| 68694-11-1 | Triflumizole | 1.81 | 1.43 | Pesticide |

